# HAMMER: Hairpin-based APOBEC3A-mediated mRNA editing reporter

**DOI:** 10.64898/2025.12.22.695965

**Authors:** Yanjun Chen, Christopher D. Mullally, Bojana Stefanovska, Reuben S. Harris

## Abstract

APOBEC3A catalyzes cytosine-to-uracil deamination in single-stranded DNA and RNA. Physiologically, APOBEC3A functions in innate immunity and aberrant deamination is associated with cytosine mutations in enzymatically preferred YTCW substrate motifs in multiple cancers. Much less is known about the potential contribution of APOBEC3A-catalyzed RNA editing to virus and cancer evolution. Here, we present HAMMER (hairpin-based APOBEC3A-mediated mRNA editing reporter), a rapid luminescence-based cellular assay for measuring RNA editing by APOBEC3A. HAMMER reports APOBEC3A activity as a reduction in the ratio of firefly to renilla luciferase activity. Briefly, tandem renilla and firefly luciferase open reading frames are separated by an optimal APOBEC3A hairpin substrate, in which C-to-U editing of a <u>C</u>GA motif yields a <u>U</u>GA stop codon thus preventing translation of the downstream firefly luciferase reporter, without impacting the upstream renilla reporter. HAMMER activation is dose-responsive, catalytic activity-dependent, and specific to human APOBEC3A. A panel of herpesviral ribonucleotide reductase constructs was used to show that direct inhibition of APOBEC3A results in a dose-responsive recovery of firefly luciferase expression. HAMMER is therefore a scalable and easy-to-use method for quantifying cellular APOBEC3A RNA editing activity and characterizing inhibitors.

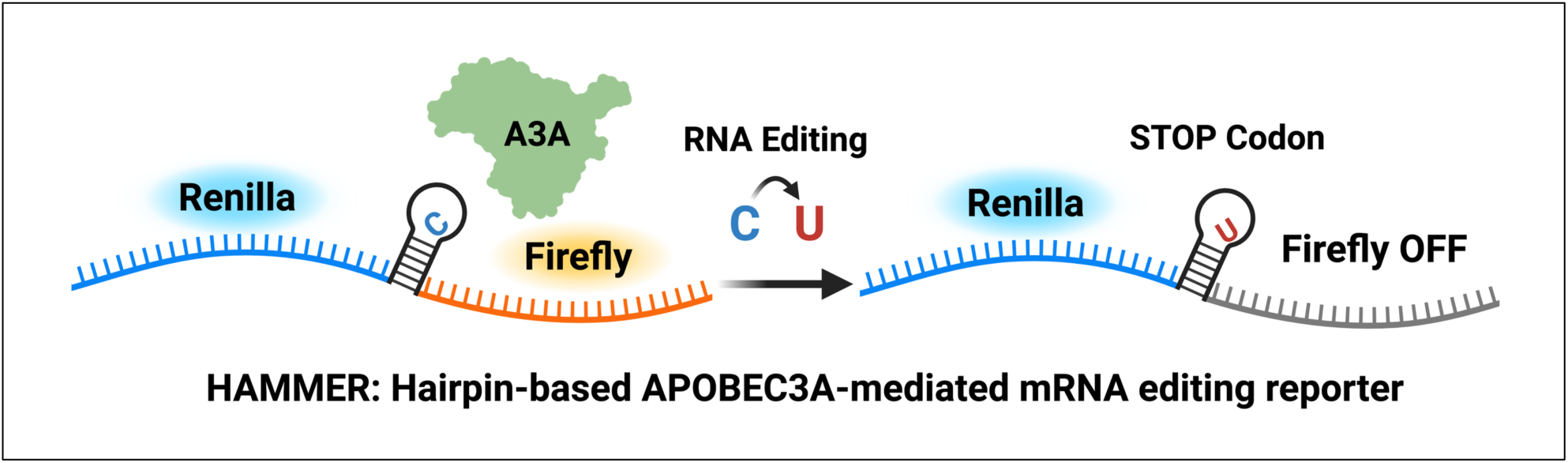

## Introduction

The seven human APOBEC3 enzymes (A3A-D and A3F-H) are generally thought to function as an overlapping innate immune barrier to endogenous and exogenous virus replication (reviewed by [1–5]). A variety of different viral families have been reported as substrates including retroviruses (HIV-1, HIV-2, HTLV), herpesviruses (EBV, KSHV, HSV-1), papillomaviruses (HPV), polyomaviruses (BKPyV, JCPyV), adenoviruses, parvoviruses, and others [1,3,6]. Key evidence for effector functions in antiviral immunity include potent virus restriction activity, copy number variability, genetic polymorphisms (positive selection), interferon-inducibility, and existence of an equally strong viral counterdefense mechanism [7]. As an example, human APOBEC3B (A3B)-mediated restriction of Epstein Barr virus (EBV) is counteracted by the large subunit of the viral ribonucleotide reductase (RNR) [8], and the copy numbers of human *A3B* and related *A3* genes vary between species [9], *A3B* itself is polymorphic within most species including humans [10–12], and *A3B* is regulated by infection and immunity signaling networks including (non)canonical NF-κB and interferon-α [13–17]. Despite sharing a common ancestor with present-day APOBEC3s, the related deaminases AID and APOBEC1 have different biological functions – AID as a genomic DNA deaminase in antibody gene diversification through somatic hypermutation and class switch recombination and APOBEC1 in mRNA editing (C6666 of the canonical *APOB* mRNA and many other coding and non-coding RNA substrates) [4].

APOBEC3A (A3A) is the most potent single-stranded (ss)DNA deaminase in humans [18–20]. A3A has a larger active site cavity than related human family members, which enables a broader range of substrates including normal cytosines in ssDNA or RNA as well as 5-methyl-C nucleobases in ssDNA [21–24]. Although A3A has been reported to restrict several of the viruses listed above, viral counterdefense mechanisms that selectively block this deaminase have yet to be described. Thus, unambiguous physiologic substrates have yet to be identified for this enzyme, although high relative activity and promiscuity are consistent with a more general role in foreign nucleic acid clearance [25,26]. The RNA cytosine deamination activity of A3A is not as well characterized as its DNA editing activity. First discovered in human macrophages during M1 polarization and in monocytes in response to hypoxia and interferons, the RNA editing activity of A3A appears capable of impacting hundreds of sites including dozens at high efficiencies [27,28]. Many of the highest efficiency A3A RNA editing sites, including *SDHB* C136, are located in loop regions of hairpin structures [29,30], which is supported by structural studies with U-shaped ssDNA and hairpin DNA structures [21,23,31]. Interestingly, A3A signature edits have been reported in SARS-CoV-2 genomic RNA sequences suggesting a potential role for this deaminase in RNA virus mutation [32–34].

In contrast to beneficial roles in innate immune function, A3A can also be pathogenic in humans by contributing to cancer mutagenesis (reviewed by [5,35,36]). The broader substrate preference of A3A and the related enzyme A3B, YT<u>C</u>W and RT<u>C</u>W, respectively, is evident in nearly 70% of human tumor genomic DNA sequences and together are the largest sources of mutation in several cancer types including bladder and cervical [36–39]. C-to-T and C-to-G single base substitution mutations in T<u>C</u>W motifs comprise SBS2 and SBS13 mutation signatures in COSMIC [37]. Ectopic expression of A3A or A3B in mice increases rates of tumor formation and leads to APOBEC signature mutations [40–42]. A3A and A3B have also been shown to cause drug resistance mutations and associate with tumor metastases [43–45]. Several strategies are being developed to diagnose and treat A3A/B-positive tumors with the most straightforward being direct inhibition of the deaminases themselves [35,46,47].

However, a major hurdle to studying A3A function and evaluating candidate inhibitors is a lack of cell-based assays. This is in part due to toxicity of long-term ectopic over-expression of wildtype human A3A. One popular assay utilizes a cytosine base editing (CBE) complex to transiently deliver A3A to a ssDNA cytosine target in an R-loop, which can be read out as a restoration of fluorescence [48–52]. Another assay leverages highly sensitive digital droplet PCR technology to quantify A3A-exclusive RNA editing hotspots, such as hairpin loop cytosines in *SDHB* or *DDOST* mRNAs [30]. Here, we report the development of HAMMER – hairpin-based A3A-mediated mRNA editing reporter – which utilizes a *DDOST*-derived hairpin substrate to provide a rapid luminescence-based readout of the RNA editing activity of A3A. The assay is highly specific to human A3A and, importantly, the luminescent read-out is directly proportional to the quantification of site-specific RNA editing by sequencing. HAMMER is therefore a robust cellular assay that reports the mRNA editing activity of human A3A. HAMMER has advantages over other systems including the fact that A3A activity strongly suppresses luminescence and then A3A inhibition is able to promote a gain-of-signal readout (not a loss-of-signal), which facilitates inhibitor testing as evidenced here by surveying a panel of candidate A3A antagonists from herpesviruses.

## Materials and Methods

### Plasmids

The HAMMER *DDOST* Hairpin1 reporter was created using In-fusion Snap Assembly (Takara, 638948) to insert a synthetic gBlock (Integrated DNA Technologies) containing a renilla luciferase open reading frame (GenBank KF035113.1), a linker harboring a modified *DDOST* hairpin sequence and multiple restriction sites, and a firefly luciferase open reading frame (GenBank MT119956.1). The linker gBlock sequence is provided in **Table S1**. Additional HAMMER reporters were also constructed using In-fusion Snap Assembly by linearizing the *DDOST* Hairpin1 reporter linker region with HindIII (New England Biolabs, R3104L) and XbaI (New England Biolabs, R0145L) and inserting gBlocks containing candidate A3A hairpin substrates (gBlock sequences in **Table S1**). Stop1 and Linear1 control reporter plasmids were generated by site-directed mutagenesis of the *DDOST* Hairpin1 reporter construct (oligonucleotide sequences in **Table S2**).

pcDNA3.1 plasmids encoding untagged human A3A and A3A-E72A as well as C-terminal Myc-His tagged human A3A, A3B, A3Bctd, A3C, A3D, A3F, A3G, A3H-HapII, AID, and APOBEC1 deaminases have been described [19,53]. pcDNA4/TO plasmids encoding C-terminal FLAG-tagged RNRs and C-terminal eGFP-tagged A3A of *Homo sapiens* (human), *Macaca mulatta* (rhesus macaque), and *Callithrix jacchus* (common marmoset) have also been reported [54,55]. GenBank accessions of the RNR constructs are described in **Table. S3**. Deaminase constructs correspond to the following GenBank accessions: *Homo sapiens* A3A (NM_145699.4), *Homo sapiens* A3B (NM_004900), *Homo sapiens* A3C (NM_014508.3), *Homo sapiens* A3D (NM_152426.4), *Homo sapiens* A3F (NM_145298.6), *Homo sapiens* A3G (NM_021822.4), *Homo sapiens* A3H-HapII (FJ376615.1); *Homo sapiens* AID (AICDA, NM_020661.4); *Homo sapiens* APOBEC1 (NM_001644.5); *Macaca mulatta* A3A (NM_001246231.1), and *Callithrix jacchus* A3A (NM_001301845.1). Expression plasmids for A1CF and RBM47 were obtained from AddGene (#112860 and #134607, respectively).

The DNA editing AMBER reporter and corresponding gRNA expression vector have been described [49]. The pCMV-Cas9n-BE4max construct was generated by site-directed mutagenesis to remove the rAPOBEC1 coding sequence [56,57]. Oligonucleotide sequences are provided in **Table S2**.

### Cell culture

293T cells (ATCC CRL-3216) were maintained in RPMI-1640 medium (Thermo Fisher Scientific) supplemented with 10% (v/v) fetal bovine serum (Biowest) and 1% (v/v) penicillin-streptomycin (Gibco), at 37 °C with 5% CO_2_.

### HAMMER assay transfection and luciferase quantification

293T cells were seeded in 12-well plates (Corning) and transfected at approximately 85% confluency. For standard experiments, 300 ng of HAMMER reporter plasmid, 200 ng of the indicated APOBEC expression plasmid, as well as 500 ng of empty vector carrier DNA were co-transfected using 3 µL TransIT-LT1 (Mirus, #MIR 2300) per well, following the manufacturer’s protocol.

At 4 hrs post-transfection, cells were trypsinized, and one-tenth of the cells from each well were transferred into a 96-well plate (Thermo Fisher Scientific, 65306) for luciferase measurements the next day. The remaining cells were replated into the original 12-well plate and collected the next day for immunoblotting and nucleic acid extraction (described below). Luciferase activity was measured 24 hrs post-transfection using the Dual-Glo Luciferase Assay System (Promega) according to the manufacturer’s instructions, and luminescence was quantified using a Spark multimode microplate reader (Tecan). Under these transient expression conditions, the firefly luminescence values of the Hairpin1 reporter are approximately 500,000 (no A3A) and the background from the Stop1 reporter is approximately 1,000-2,000, yielding a theoretical signal:noise ratio of approximately 250– to 500-fold. Though active human A3A yields a robust and dose-responsive signal in this system, these maximal values are not reached due to incomplete editing in cells expressing A3A and some cells potentially only receiving Hairpin1 plasmid with no/low A3A.

### Sanger sequencing of total DNA samples

For DNA analysis, transfected cells reseeded in 12-well plates were harvested at 24 hrs post-transfection. One-tenth of the cells from each well was pelleted and resuspended in QuickExtract DNA Extraction Solution (Biosearch Technologies, SS000035-D2) according to the manufacturer’s protocol to obtain PCR-ready total DNA solution. 1 µL of processed solution was used as template for PCR amplification of the HAMMER reporter linker region using primers listed in **Table S2**. PCR products were size verified on a 1% agarose gel and purified using the Monarch PCR & DNA Cleanup Kit (NEB, T1135). Purified amplicons were Sanger sequenced by Eurofins using primers listed in **Table S2**. Chromatograms received from Sanger sequencing were uploaded to EditR for peak quantification [58].

### Reverse transcription and Sanger sequencing of cDNA samples

For RNA analysis, transfected cells in 12-well plates were harvested at 24 hrs post-transfection, and one-tenth of the cells from each well was pelleted for total RNA purification using the RNeasy Mini Kit (Qiagen, 74104). One microgram of purified RNA was reverse-transcribed using Transcriptor First Strand cDNA Synthesis Kit (Roche, #04897030001) according to the manufacturer’s instructions. Purified cDNA was used as template for PCR amplification of the HAMMER reporter linker region using primers listed in **Table S2**. PCR products were purified, sequenced, and quantified as described above.

### Immunoblotting

For protein analysis, transfected cells in 12-well plates were harvested 24 hrs post-transfection, and about 80% of the cells remained after collection for DNA and RNA analysis described above. Cells were PBS washed, lysed in RIPA Buffer (Thermo Fisher Scientific, PI89900), and quantified by BCA assay (Thermo Fisher Scientific, 23225). Samples were then separated by 4-20% tris glycine polyacrylamide gels (Bio-Rad, 5671095) run for 90 minutes at 120V and transferred to PVDF-FL membrane (Millipore Sigma, IPFL00005). Following transfer, membranes were washed with PBS 0.1% Tween (PBST) before incubation in 1x casein blocking buffer (Sigma-Aldrich, C7594). Primary antibodies were diluted in 1x casein blocking buffer and incubated on membrane overnight at 4°C. Membranes were washed 3 times in PBST and incubated with secondary antibodies diluted in 1x blocking buffer supplemented with 0.01% SDS. Following 1 hr of incubation at room temperature and 3 rounds of washing with PBST, membranes were imaged on LiCor Odyssey Imager. Primary antibodies used were anti-A3A (5210-87-13 [59]; 1:1,000), anti-GFP (Cell Signal Technology, 2555S; 1:1,000), anti-Myc (Cell Signal Technology, 2278S; 1:1,000), anti-Flag (Sigma-Aldrich, F1804; 1:1,000), anti-Renilla luciferase (Abcam, EPR17792; 1:1,000), anti-β-Actin (Sigma Aldrich, A1978; 1:5,000), and anti-Tubulin (Sigma Aldrich, T5168; 1:5,000); Secondary antibodies used were anti-rabbit IRdye 800CW (LI-COR, 827-08365; 1:10,000) and anti-mouse IRdye 680LT (LI-COR, 925-68020; 1;10,000).

### HAMMER assays to test RNR inhibitory activity

293T cells were seeded and transfected as described above. Specifically, cells were co-transfected with 300 ng HAMMER reporter plasmid, 200 ng A3A expression plasmid, 400 ng of the indicated RNR expression plasmid, and 100 ng empty vector carrier DNA using 3 µL TransIT-LT1 per well. Sample processing and luciferase measurements were performed identically to the standard HAMMER workflow.

### AMBER assays to test RNR inhibitory activity

293T cells were seeded in 12-well plates and transfected as described above. Cells were co-transfected with 300 ng of AMBER reporter plasmid, 200 ng of Cas9n-BE4max plasmid, 100 ng of AMBER gRNA plasmid, 200 ng of A3A expression plasmid, and 400 ng of the indicated RNR expression plasmid using 3.6 µL TransIT-LT1 per well. At 24 hrs, cells were trypsinized and suspended in 1 mL of PBS. An aliquot of 200 µL was used for flow cytometry to quantify mCherry and eGFP expression, and the remaining cells were pelleted for immunoblot analysis as described above.

### Phylogenetic analysis of viral RNRs

Viral RNR amino-acid sequences (GenBank accessions listed in **Table S3**) were assembled in FASTA format and aligned with MAFFT (v7.490) [60]. The alignment was trimmed with trimAl (v1.4) to reduce poorly aligned and gap-rich regions. A maximum-likelihood phylogeny was inferred from the trimmed alignment using FastTree (v2.1.11) under the Jones–Taylor–Thornton (JTT) substitution model [61]. The resulting tree was midpoint-rooted (phytools in R) and visualized in R (v4.5.0) using ggtree (v4.0.1) [62].

### Data analysis and figure generation

Illustrations of reporter constructs were generated with BioRender. Data analysis and figure generation were performed in GraphPad Prism. Reporter activation is calculated as the ratio of luminescence, Firefly/Renilla, and normalized to empty vector or catalytic mutant control where indicated. The student’s t-test was used to assess differences in reporter activation, and Pearson correlation was used to determine correlation between reporter activation and DNA/RNA editing. A threshold for significance was set at *P* < 0.05. Samples showing statistically significant differences relative to the indicated controls are annotated with *P*-values, and unlabeled comparisons did not reach the significance threshold.

## Results

### Construction of HAMMER – <u>h</u>airpin-based <u>A</u>POBEC3A-<u>m</u>ediated <u>m</u>RNA <u>e</u>diting reporter

The catalytic activity of A3A is strongly influenced by nucleic acid secondary structure. For instance, A3A exhibits a pronounced preference for cytosine nucleobases positioned on the 3’ side of RNA and DNA hairpin loops [22,29,30,63–66]. This preference inspired the design of a series of dual-luciferase reporter plasmids to express in-frame renilla and firefly luciferase genes separated by a short linker sequence containing an editable hairpin (**Fig. 1A**). The initial Hairpin1 construct was based upon a naturally occurring hairpin in the *DDOST* transcript, because deamination of *DDOST* hairpin loop C558-to-U has been reported as a biomarker of A3A expression and activity [30]. However, in order to convert this hairpin into a viable reporter, the native *DDOST* hairpin had to be altered by swapping a single A-U base pair into U-A within the top of the hairpin stem (Hairpin1 in **Fig. 1A**). This change enabled a single C-to-U editing event within the hairpin loop <u>C</u>GA motif to yield a <u>U</u>GA stop codon, which completely blocks translation of the downstream firefly gene but does not alter translation of the upstream renilla gene (**Fig. 1A**). Thus, A3A-mediated deamination may be quantified by measuring the firefly-to-renilla luminescence ratio.

**Fig. 1.**
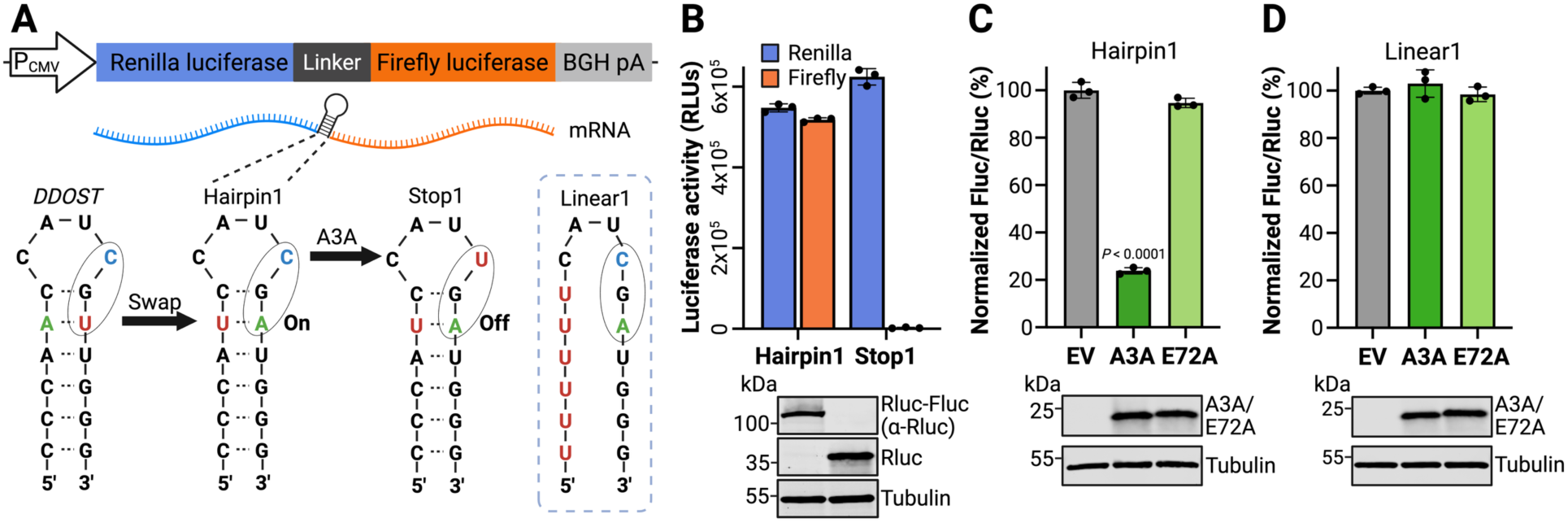
Design and validation of the HAMMER reporter. **A**, Schematic of the reporter construct with a CMV promoter, an upstream renilla luciferase (Rluc) gene, a linker region, and a downstream firefly luciferase (Fluc) gene. The linker region was engineered to contain a *DDOST-derived* hairpin (Hairpin1), an otherwise identical hairpin with a constitutive stop codon (Stop1), or a linear control sequence (Linear1). **B**, Luciferase activities of Hairpin1 and Stop1 reporters following transfection into 293T cells (RLU, relative luminescent units; mean ± SD of 3 biological replicates). Immunoblots shown below confirm expression of Rluc-FLuc chimeric protein by Hairpin1 and Rluc only by Stop1 with tubulin as a loading control. **C-D**, Normalized firefly-to-renilla luminescence ratios for Hairpin1 or Linear1 reporters co-transfected in 293T cells with an empty vector (EV), human A3A, or the human A3A-E72A catalytic mutant (mean ± SD of 3 biological replicates). Immunoblots shown below confirm expression of A3A and A3A-E72A with tubulin as a loading control.

To test the signal-to-noise ratio of the Hairpin1 reporter, 293T cells were transfected with the Hairpin1 reporter in parallel with a Stop1 control, in which the target cytosine nucleobase in the loop is replaced with uracil to encode a constitutive stop codon (**Fig. 1A**). 293T cells were selected because they lack endogenous *A3A* expression and support high transient transfection frequencies [59,67]. As expected, the Hairpin1 reporter expressed both renilla and firefly luciferase, whereas the Stop1 construct only expressed renilla luciferase activity with firefly luciferase completely suppressed by the engineered stop codon (**Fig. 1B**). Next, we tested whether the Hairpin1 reporter could detect A3A editing activity. Co-transfection of 293T cells with the Hairpin1 reporter and plasmids expressing wildtype A3A, catalytic mutant A3A-E72A, or an empty vector control showed that wildtype A3A reduces the firefly-to-renilla ratio by 80% (**Fig.1C**). In contrast, A3A-E72A had no significant effect (less than 3%). These results indicated that the Hairpin1 assay provides a direct read-out of A3A catalytic activity.

To ask whether the hairpin structure is required for reporter function, a linear control construct was generated by changing the 5’ side of the stem to uracil (Linear1 in **Fig.1A**). Importantly, the Linear1 construct preserves the UCG deamination target site, as well as two additional ribonucleobases upstream and all bases downstream. However, co-transfection of 293T cells with the Linear1 reporter and plasmids expressing wildtype A3A, catalytic mutant A3A-E72A, or an empty vector control showed that wildtype A3A does not significantly alter the expression levels of the Linear1 reporter (**Fig. 1D**). Taken together, these results combined to indicate that the A3A-dependent diminution of firefly luciferase activity requires a hairpin structure as a substrate.

In addition to *DDOST1*, several other hairpin substrates have been reported for A3A [29,30,64] (**Fig. S1A**). Dual-luciferase reporter plasmids were designed for several of these hairpin substrates, such that single C-to-U editing events would yield a stop codon. To compare with the *DDOST*-derived Hairpin1 substrate, each of these constructs was co-transfected with A3A or A3A-E72A expression plasmids. Remarkably, the *DDOST*-derived Hairpin1 substrate was found to support the highest level of editing activity by A3A (**Fig. S1B**). *CYFIP1* C3222– and *SDHB* C136-based hairpin constructs also supported clear A3A editing activity, though both were significantly less than that observed with the *DDOST*-derived Hairpin1.

Although the enzymology of A3A on RNA hairpins has yet to be characterized comprehensively, studies on DNA hairpins have shown that A3A prefers substrates with longer stems, smaller loops (3-nt > 4-nt), and specific contexts (TTC loop over others) [63–66]. Guided by these principles, we engineered variants of the *DDOST* Hairpin1 with altered loop sizes, loop sequences, and extended stems (**Fig. S1C**). Contrary to expectations, all modifications reduced reporter activity and the original *DDOST-*based Hairpin1 substrate remained the most effective configuration (**Fig. S1D**). Given superior performance, the *DDOST* Hairpin1 reporter was used for all subsequent experiments, and the dual luciferase reporter system incorporating Hairpin1 is hereafter called the <u>h</u>airpin-based <u>A</u>3A-<u>m</u>ediated <u>m</u>RNA <u>e</u>diting reporter (HAMMER). This reporter also supported A3A-dependent RNA editing in HeLa cells, indicating broader mechanistic functionality and utility (**Fig. S2**).

### HAMMER reports A3A mRNA editing activity dose-responsively

We next evaluated whether HAMMER could report A3A activity in a dose-responsive manner. 293T cells were co-transfected with the HAMMER reporter and varying amounts of A3A or A3A-E72A expression constructs, and luciferase activity was measured 24 hrs post-transfection. Relative to the empty vector control, wildtype A3A decreased the firefly-to-renilla luminescence ratio in a clear dose-dependent fashion, and A3A-E72A did not produce a significant change in signal, even though the wildtype and catalytic mutant proteins were expressed at similar levels in immunoblots (**Fig. 2A**). These data demonstrated that HAMMER quantitatively reports A3A activity across a wide range of expression levels.

**Fig. 2.**
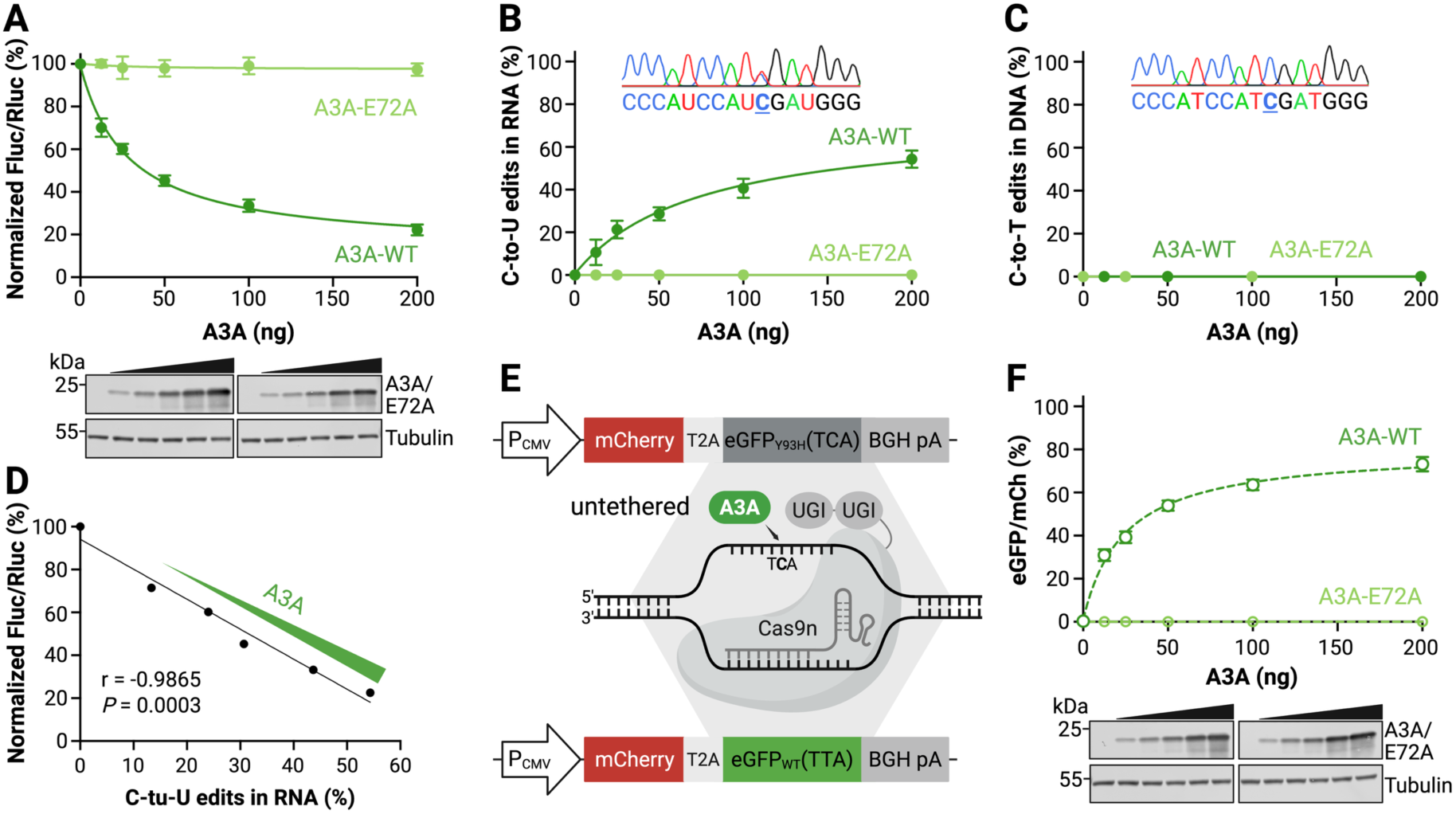
HAMMER reports A3A activity dose-dependently and specifically on RNA. **A**, Normalized firefly-to-renilla luminescence ratios of 293T cells co-transfected with HAMMER reporter and increasing amounts of A3A or A3A-E72A plasmid. Data represent mean ± SD of 3 biological replicates. Immunoblots shown below confirm dose-dependent expression of A3A and A3A-E72A. **B**, Percentage of C-to-U edits in HAMMER reporter RNA following reverse transcription, PCR amplification, and Sanger sequencing. Data represent mean ± SD of 3 biological replicates. A representative chromatogram from the 200 ng A3A expression condition is shown in the inset. **C**, Percentage of C-to-T mutations detected in reporter DNA PCR-amplified and Sanger-sequenced from the same reactions as panel B. Data represent mean ± SD of 3 biological replicates, with some error bars smaller than the data points. A representative chromatogram from the 200 ng A3A expression condition is shown in the inset. **D**, Relationship between HAMMER luminescent readout and the percent mRNA C-to-U editing by A3A (2-fold dilution series starting at 200 ng; Pearson correlation coefficient (r) and *P*-value are indicated). **E**, Schematic of the untethered DNA editing assay AMBER. An R-loop substrate created in the *eGFP* gene using a gRNA/Cas9nickase-UGI complex can be edited by A3A to restore fluorescence. **F**, eGFP fluorescence of 293T cells co-transfected with the AMBER reporter and increasing amounts of A3A or A3A-E72A plasmid. Data represent mean ± SD of 3 biological replicates, with some error bars smaller than the data points. Immunoblots shown below confirm dose-dependent expression of A3A and A3A-E72A.

As demonstrated previously, single-stranded DNA is the preferred, and often the exclusive, substrate of the seven human APOBEC3 family members [18,20,24,68]. Therefore, it is critical to determine whether the HAMMER signal originates from RNA editing rather than DNA deamination of the plasmid reporter. To directly address this possibility, total RNA and DNA were collected in parallel to the HAMMER assay. Sequencing results indicated that only the *DDOST*-derived Hairpin1 mRNA is edited by A3A, and that the corresponding DNA region is unaffected (**Fig. 2B-C**). Moreover, the luminescent readout of HAMMER (Fluc/Rluc) is directly proportional to the quantification of the C-to-U RNA editing frequencies by sequencing (**Fig. 2D**). These results confirm that HAMMER specifically reports RNA editing by A3A and is unlikely to be confounded by DNA deamination events.

As an additional control, parallel experiments were done in 293T cells using the same A3A expression constructs and a DNA C-to-T mutation reporter called AMBER (APOBEC-mediated base editing reporter) [48,49]. In this system, untethered A3A can access the target DNA cytosine in eGFP after exposure by a gRNA/Cas9nickase directed R-loop formation (assay schematic in **Fig. 2E**). Additionally, uracil excision repair is prevented by tethering two uracil DNA glycosylase inhibitory (Ugi) peptides to the C-terminal end of the Cas9nickase. As above, increasing amounts of transfected A3A but not A3A-E72A cause a dose-responsive change in reporter activity (here, increasing eGFP fluorescence; **Fig. 2F**). However, in contrast to HAMMER, all AMBER reporter activity is attributable to direct deamination of reporter DNA as established in the original reports [48,49].

### HAMMER selectively detects human A3A RNA editing activity

Despite extensive structural homology and broadly conserved enzymatic function, APOBEC enzymes exhibit marked differences in catalytic activity, substrate preference, and biological function (summarized in Introduction and reviewed in [1–4]). All human *APOBEC3* family genes were derived from ancestral *AID* and *APOBEC1* sequences by multiple gene duplication and diversification events [11,69] (schematic of phylogenetic relationships in **Fig. 3A**). All nine of these human enzymes have shown DNA deamination activity in variety of contexts, but only APOBEC1, A3A, A3B, and A3G have been reported to edit RNA [27–29,70–75].

**Fig. 3.**
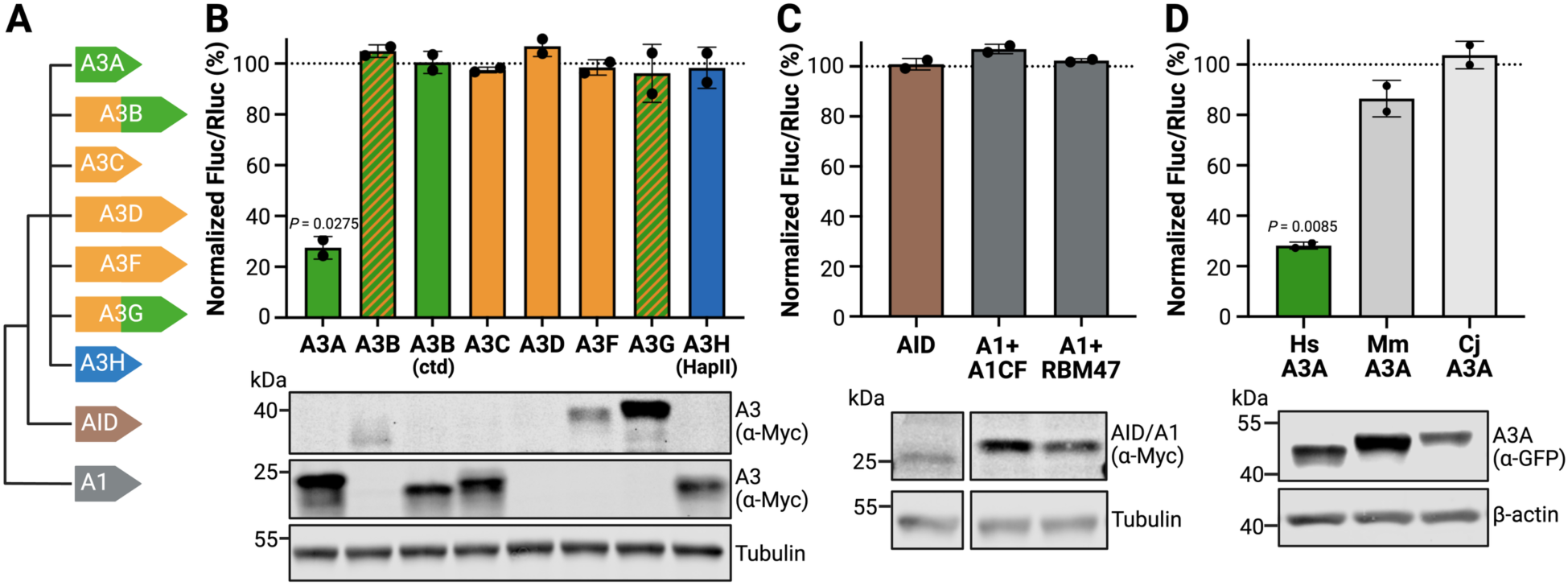
HAMMER selectively detects RNA editing by human A3A. **A**, Schematic of the nine catalytically active human APOBEC deaminase family members. Phy-logenetically distinct APOBEC3 Z1-type deaminase domains are green, Z2-types are orange, and the single Z3-type is blue. AID and A1 are indicated in different colors. **B**, Firefly-to-renilla luminescence ratios for HAMMER performed with the indicated human APO-BEC family member constructs (mean ± SD of 2 biological reactions normalized to vector con-trol). Immunoblot of APOBEC expression (anti-Myc) with tubulin as a loading control. **C**, Firefly-to-renilla luminescence ratios for HAMMER performed with AID, APOBEC1/A1CF, or APOBEC1/RBM47 (mean ± SD of 2 biological reactions normalized to vector control). Im-munoblot of APOBEC expression (anti-Myc) with tubulin as a loading control (bands are from the same blot and reorganized for presentation to eliminate irrelevant lanes). **D**, Firefly-to-renilla luminescence ratios for HAMMER performed with A3A homologs from *Homo sapiens* (Hs), *Macaca mulatta* (Mm), and *Callithrix jacchus* (Cj) (mean ± SD of 2 biological reac-tions normalized to vector control). Immunoblot of deaminase expression (anti-GFP) with β-actin as a loading control.

We therefore performed a head-to-head comparison of the nine active human APOBEC family members using the *DDOST*-derived HAMMER assay with the Hairpin1 substrate. First, 293T cells were co-transfected with HAMMER and expression plasmids encoding A3A, A3B, A3C, A3D, A3F, A3G, and A3H (**Fig. 3B**). Among >10 A3H haplotypes, HapII was used here because it is a common allele in the human population and the encoded enzyme is stable and catalytically active [76,77]. The C-terminal catalytic domain of A3B (A3Bctd) was also tested because it shares 89% amino acid identity (93% similarity) with A3A. As above, renilla and firefly luciferase activity was measured 24 hrs post-transfection. Among these constructs, only A3A edited the reporter as evidenced by a marked decrease in the firefly-to-renilla luminescence ratio (**Fig. 3B**). Even A3Bctd, which expressed to nearly the same extent as A3A, showed no detectable RNA editing activity. The double-domain enzymes, A3B, A3D, A3F, and A3G were not expressed as well as the single domain deaminases A3A, A3C, and A3H-Hap II and, despite repeated co-transfection experiments, these constructs showed no signs of activity in the HAMMER assay. This negative result for A3D was reconfirmed in a dose-responsive reconstruction experiment (**Fig. S3**). Additional experiments also showed that neither human AID nor human APOBEC1 exhibit activity in HAMMER, even when the latter deaminase was co-expressed with physiologically relevant co-factors A1CF or RBM47 [78–80] (**Fig. 3C**). These results combined to indicate that HAMMER selectively reports human A3A activity.

As an additional test of specificity, HAMMER activity was compared for human A3A and homologs from two non-human primate species – the Old World rhesus macaque (*Macaca mulatta*) and the New World common marmoset (*Callithrix jacchus*). These A3A proteins are 81% (88%) and 71% (82%) identical (similar) to the human enzyme. Interestingly, although human A3A exhibited the strongest activity in HAMMER, the rhesus macaque A3A protein showed a modest effect, and the common marmoset A3A exhibited little to no activity (**Fig. 3D**). These experiments combine to underscore the specificity of the HAMMER assay for human A3A, because even some of its closest naturally occurring homologs (A3Bctd, MmA3A, and CjA3A) are unable to approach its robust Hairpin1 RNA editing activity.

### HAMMER identifies herpesvirus RNRs that inhibit A3A

Prior studies have shown that the EBV large RNR subunit BORF2 directly binds to A3Bctd, inhibits its catalytic activity, and triggers its accumulation into perinuclear aggregates [8,81,82]. As noted above, human A3A is 89% identical/93% similar to human A3Bctd, and it can also be bound by EBV BORF2 and relocalized into aggregates [54,82]. Therefore, as a first proof-of-concept, we sought to test whether EBV BORF2 and related herpesviral RNR large subunits have the capacity to inhibit A3A-catalyzed RNA editing using HAMMER. For comparison, all RNR constructs were also tested in parallel for inhibition of A3A-catalyzed DNA editing using an untethered version of AMBER (schematic in **Fig. 2E**). The herpesvirus large RNR subunits surveyed here are derived from the human α-herpesvirus HSV-1 and from human and Old and New World primate γ-herpesviruses from the lymphocryptovirus and rhadinoherpesvirus groups (phylogenetic relationships in **Fig. 4A**).

**Fig. 4.**
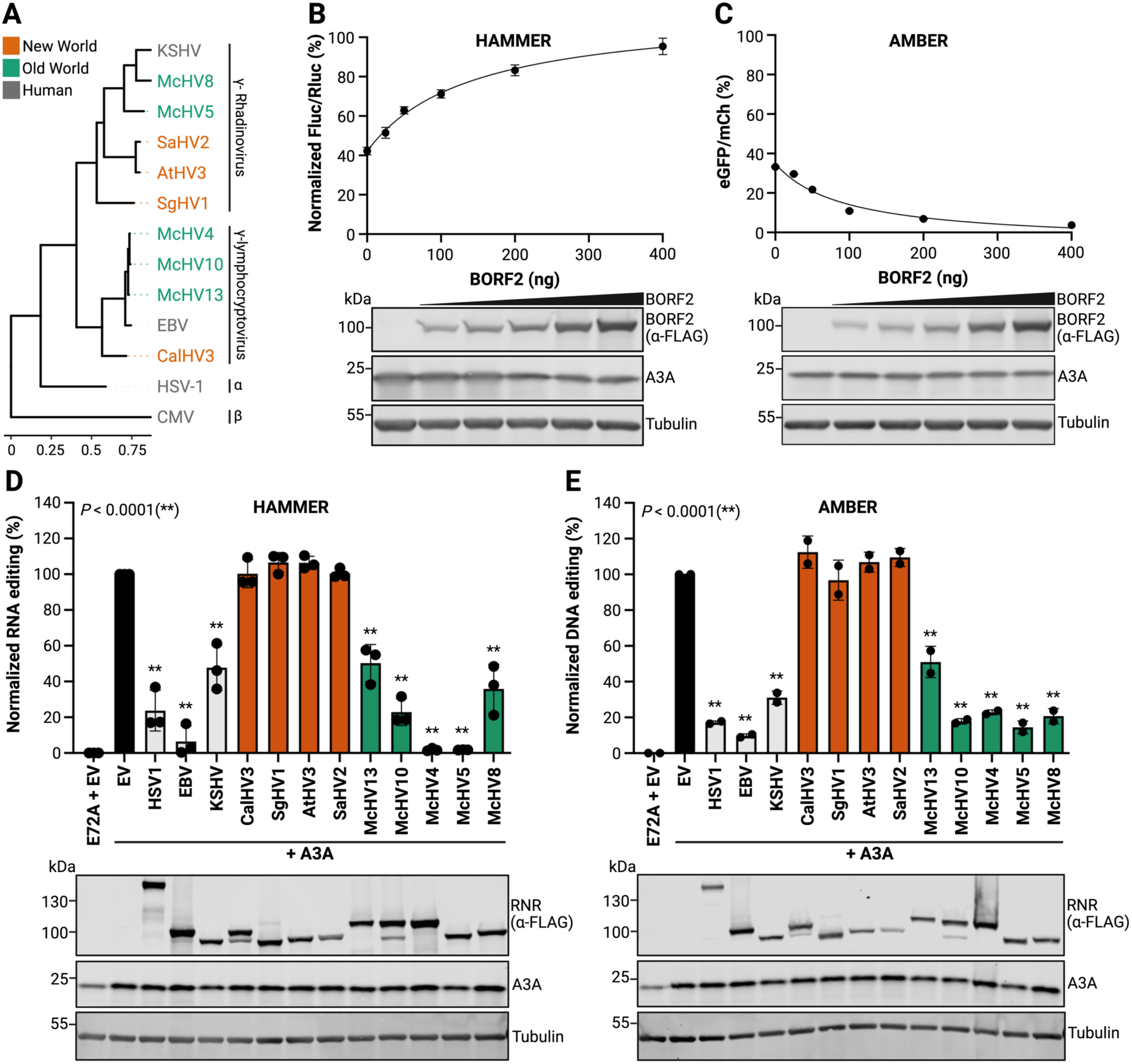
HAMMER identifies herpes virus RNRs that inhibit A3A activity. **A**, Phylogenetic tree of viral RNR family members tested in HAMMER and AMBER assays (the β-herpesvirus human CMV is included as a more distantly related outgroup). **B-C**, HAMMER and AMBER results, respectively, for 293T cells co-transfected with each reporter, 200 ng A3A, and the indicated amounts of an EBV BORF2 construct. Data represent the mean ± SD of technical triplicate experiments. Immunoblots showing A3A and BORF2 (anti-FLAG) expression with tubulin as a loading control. **D-E**, HAMMER and AMBER results, respectively, for 293T cells co-transfected with each reporter, 200 ng A3A, and the indicated viral RNR constructs. HAMMER activity was quantified by normalizing firefly-to-renilla luminescence ratios to EV control (100%) and E72A control (0%). AMBER activity was quantified by normalizing eGFP+ to mCherry+ cell count ratios to EV control (100%) and E72A control (as 0%). Data represent mean ± SD of three (HAMMER) or two (AMBER) biological experiments. Immunoblots show A3A and RNR (anti-FLAG) expression with tubulin as a loading control.

First, titration experiments were performed to ask whether BORF2 inhibits A3A-catalyzed HAMMER read-out dose-responsively. As above, A3A caused a 60% reduction in firefly luciferase in HAMMER and, interestingly, BORF2 restored luminescence dose-responsively to near wildtype levels at the highest transfected amounts (**Fig. 4B**). In a parallel experiment, A3A caused a 30% restoration of eGFP fluorescence in AMBER, and BORF2 inhibited this DNA editing activity dose responsively with near-complete inhibition at highest expression levels, consistent with prior reports [49,83] (**Fig. 4C**). A strong correlation between RNA and DNA editing inhibition indicated that EBV BORF2 is capable of inhibiting both of these human A3A activities (**Fig. S4**).

Second, the large RNR subunit of HSV-1 (ICP6) was tested in comparison with that of lymphocryptovirus EBV (BORF2) and the rhadinoherpesvirus KSHV (ORF61). Relative to EBV BORF2, which inhibited A3A strongly, the large RNR subunits of HSV-1 and KSHV appeared to have intermediate inhibitory activities (**Fig. 4D-E**). Third, RNRs from several New World primates were tested and none of these proteins appeared to inhibit either the RNA or the DNA editing activity of human A3A (**Fig. 4D-E**). Fourth and in striking contrast to the New World viral RNRs, the large RNR subunits from several Old World primates were tested and, surprisingly, all of them inhibited both RNA and DNA editing activity of human A3A (**Fig. 4D-E**). Most remarkably, two of these Old World viral RNRs, the large RNR subunits from the lymphocryptovirus McHV4 and the rhadinoherpesvirus McHV5 proved to be as potent as EBV BORF2 at blocking A3A activity (**Fig. 4D-E**). These results demonstrate that HAMMER can be used to identify candidate A3A inhibitors by virtue of clear gain-of-signal readouts.

## Discussion

Here, we present HAMMER (hairpin-based APOBEC3A-mediated mRNA editing reporter) for quantification of the RNA editing activity of human A3A in living cells. A3A dose-responsively edits a *DDOST*-derived hairpin substrate (Hairpin1), which yields a stop codon preventing expression of the downstream firefly luciferase reporter and leaving the upstream renilla luciferase reporter unaffected. Thus, RNA editing by A3A becomes a simple metric of the ratio of firefly to renilla luminescence values. Importantly, sequencing showed that human A3A specifically edits the reporter mRNA and does not significantly alter the corresponding DNA sequence in the transfected HAMMER construct. HAMMER is highly specific to RNA editing by human A3A, because no other active human polynucleotide deaminase exhibits activity in this assay. Even human A3Bctd (93% similar to human A3A) and the textbook RNA editing enzyme APOBEC1 show no activity in HAMMER. However, the 88% similar rhesus macaque A3A protein exhibits modest activity in HAMMER suggesting that relatively small amino acid and structural differences are likely to account for the optimal activity of human A3A with this reporter system.

How does HAMMER compare to previously developed RNA editing reporters? As far as we’re aware, all prior cellular reporters are fluorescence-based and not luminescence-based like HAMMER. One reporter has 300 bp of *APOB* sequence situated in-frame between an mCherry control marker and a downstream eGFP reporter [84]. As *APOB* is a primary physiological substrate of APOBEC1, mRNA editing by this deaminase yields a stop codon and lower eGFP levels. Though not tested against other deaminases, it is likely that this assay is specific to APOBEC1 because it includes *cis*-mooring sequences required for mRNA editing by this enzyme. Two additional assays utilize eGFP subcellular localization as a biomarker for mRNA editing [80,85,86]. One of these assays placed an mRNA editing substrate in-frame between an upstream eGFP marker and a downstream nuclear export sequence (NES) [80,86]. The other situated an *APOB* editing target region, as above, between eGFP and a cytoplasmic transmembrane targeting motif [85]. In both instances, editing creates a stop codon that uncouples eGFP from the subcellular localization determinants and results in a redistribution of eGFP reporter from the cytoplasm to the whole cell due to its relatively small size and capacity to diffuse through nuclear pores. *DDOST* or *DDOST*-like hairpin sequences were not tested in either assay, though significant mRNA editing by A3A could be detected for the related *SDHB* hairpin and an unrelated *EVI2B* hairpin in the NES-dependent assay. Like HAMMER with the *DDOST*-derived hairpin1 substrate, the *EVI2B* version of the NES-dependent assay was not edited by other APOBEC family members. However, unlike HAMMER, these relocalization-based assays have relatively modest signal-to-noise ratios and are unlikely to be scalable for high-throughput identification of A3A inhibitors.

We anticipate that HAMMER will become a valuable addition to the toolbox of cellular assays used to study A3A function in biology and disease [87]. As a cell-based RNA-specific assay for A3A deaminase activity, HAMMER will complement DNA-based assays such as AMBER used here and others that similarly leverage tethered or untethered cytosine base editing technologies [48–52,88,89]. As outlined in Introduction, A3A has been reported to interact with a broad number of DNA and RNA-based viruses. However, a true physiological substrate for A3A remains debatable because a key hallmark of a virus restriction factor has yet to be satisfied – *i.e*., a virus has yet to be found that encodes an equally potent A3A neutralization mechanism. It is therefore possible that HAMMER could be used to screen for such a factor and, minimally, once discovered to help characterize the underlying molecular mechanism. Here, as proof-of-concept, we tested a panel of herpesviral RNR large subunits against human A3A. Although human and Old World monkey A3B enzymes are clearly physiological targets of several of these viral RNRs [54], the high degree of similarity between the C-terminal catalytic domain of human A3B and human A3A suggested that some of the viral RNRs might also inhibit A3A through molecular mimicry [54]. Indeed, in addition to the RNR large subunits from several human viruses (EBV, KSHV, and HSV-1), several Old World primate viral RNR large subunits were also able to inhibit human A3A activity in HAMMER. Notably, the RNR large subunits from McHV4 and McHV5 were as potent as EBV BORF2 at blocking the mRNA editing activity of human A3A. Likewise, HAMMER is also envisaged to be useful for identifying and characterizing small molecules that impact A3A function in cells.

## Acknowledgments

We are grateful to Michael Carpenter, Allen York, and other lab members for thoughtful feedback.

R.S.H. is an Investigator of the Howard Hughes Medical Institute and the Ewing Halsell President’s Council Distinguished Chair at University of Texas San Antonio.

## Supplementary Data

Supplementary Data are available at NAR Online.

## Data Availability

All data are reported in the main manuscript or supplementary material.

## Author Contributions

Conceptualization: All authors. Methodology: YC, CM, RSH. Investigation: YC, BS, CM. Visualization: All authors. Funding acquisition: All authors. Simulations and Data Analysis: All authors. Project administration: RSH. Supervision: RSH. Writing – original draft: All authors. Writing – review & editing: All authors.

