## Supplementary Tables S1-S3 and Figures S1-S4 for "HAMMER: Hairpin-based APOBEC3A-mediated mRNA editing reporter"

**Supplementary Figures S1-S4**

### Table S1. gBlocks.

| Name | Sequence 5' to 3' |
|---|---|
| *DDOST* hairpin1 linker | AAGCTTCGTGGCAGACACCGAGAACCTGCTGAAGGCCCCATCCATCGATGGGAAATAAAGTGTAAAGAACATACTCTAGAGGGCCCA |
| *DDOST* hairpin2 gBlock | GAGCGCGTGCTGAAGAACGAGCAGAAGCTTCGTGGCAGACACCGAGAACCTGCTGAAGGCACCCATCATCGATGGGAAATAAAGTGTAAAGAACATACTCTAGAGGGCCCAGAAGACGCCAAAAACATA |
| *DDOST* hairpin3 gBlock | GAGCGCGTGCTGAAGAACGAGCAGAAGCTTCGTGGCAGACACCGAGAACCTGCTGAAGGCACCCATCTTCGATGGGAAATAAAGTGTAAAGAACATACTCTAGAGGGCCCAGAAGACGCCAAAAACATA |
| *DDOST* hairpin4 gBlock | GAGCGCGTGCTGAAGAACGAGCAGAAGCTTCGTGGCAGACACCGAGAACCTGGTTTAGGCCCCATCCATCGATGGGGCCTAAAGTGTAAAGAACATACTCTAGAGGGCCCAGAAGACGCCAAAAACATA |
| *DDOST* hairpin5 gBlock | GAGCGCGTGCTGAAGAACGAGCAGAAGCTTCGTGGCAGACACCGAGAACCTGGATTTAGGCCCCATCATCGATGGGGCCTAAAGTGTAAAGAACATACTCTAGAGGGCCCAGAAGACGCCAAAAACATA |
| *DDOST* hairpin6 gBlock | GAGCGCGTGCTGAAGAACGAGCAGAAGCTTCGTGGCAGACACCGAGAACCTGGATTTAGGCCCCATCTTCGATGGGGCCTAAAGTGTAAAGAACATACTCTAGAGGGCCCAGAAGACGCCAAAAACATA |
| *DDOST* hairpin7 gBlock | GAGCGCGTGCTGAAGAACGAGCAGAAGCTTCGTGGCAGACACCGAGAACTTGGTTTAGGCCCCATCCATCGATGGGGCCTAAACCAAAAAGAACATACTCTAGAGGGCCCAGAAGACGCCAAAAACATA |
| *DDOST* hairpin8 gBlock | GAGCGCGTGCTGAAGAACGAGCAGAAGCTTCGTGGCAGACACCGAGAAACTTGGTTTAGGCCCCATCATCGATGGGGCCTAAACCAAAAAGAACATACTCTAGAGGGCCCAGAAGACGCCAAAAACATA |
| *DDOST* hairpin9 gBlock | GAGCGCGTGCTGAAGAACGAGCAGAAGCTTCGTGGCAGACACCGAGAAACTTGGTTTAGGCCCCATCTTCGATGGGGCCTAAACCAAAAAGAACATACTCTAGAGGGCCCAGAAGACGCCAAAAACATA |
| *CYFIP1* hairpin gBlock | GAGCGCGTGCTGAAGAACGAGCAGAAGCTTCGTGGCAGACACCGAGAACCTGCTGAAGGAAATTTCCATCGAAAGAGAATAAAGTGTAAAGAACATACTCTAGAGGGCCCAGAAGACGCCAAAAACATA |
| *SDHB* hairpin gBlock | GAGCGCGTGCTGAAGAACGAGCAGAAGCTTCGTGGCAGACACCGAGAACCTGCTGAAGGCACCATCTATCGATGGGAAATAAAGTGTAAAGAACATACTCTAGAGGGCCCAGAAGACGCCAAAAACATA |
| *NUP93* hairpin gBlock | GAGCGCGTGCTGAAGAACGAGCAGAAGCTTCGTGGCAGACACCGAGAACCTGCTGATCAGCAAGCTCATCAGCTTGCTGTAAAGTGTAAAGAACATACTCTAGAGGGCCCAGAAGACGCCAAAAACATA |
| *MD21D2* hairpin gBlock | GAGCGCGTGCTGAAGAACGAGCAGAAGCTTCGTGGCAGACACCGAGAACCTGCTGAATTGCAGGCCTATCAGGCCTGCATAAAGTGTAAAGAACATACTCTAGAGGGCCCAGAAGACGCCAAAAACATA |
| *FAM83G* hairpin gBlock | GAGCGCGTGCTGAAGAACGAGCAGAAGCTTCGTGGCAGACACCGAGAACCTGCTGAATCGGGCCCCTCTCAGGGGCCCGTAAAGTGTAAAGAACATACTCTAGAGGGCCCAGAAGACGCCAAAAACATA |

**Table S2. Oligonucleotide sequences.**

| Oligo name | Purpose | Sequence 5' to 3' |
|---|---|---|
| Linear1-F | Site-directed mutagenesis for making *DDOST* Linear1 reporter (forward) | CTGAAGGCTTTTTTCATCGATGGGAAATAAAG TGT |
| Linear1-R | Site-directed mutagenesis for making *DDOST* Linear1 reporter (reverse) | CATCGATGAAAAAAGCCTTCAGCAGGTTCTCG |
| Stop1-F | Site-directed mutagenesis for making *DDOST* Stop1 reporter (forward) | CCCATCCATTGATGGGAAATAAAGTGT |
| Stop1-R | Site-directed mutagenesis for making *DDOST* Stop1 reporter (reverse) | CATCAATGGATGGGGCCTTCAGCA |
| Cas9n-BE4max-F | Site-directed mutagenesis for removal of rAPOBEC1 from BE4max (forward) | GGAAAGTCGACAAGAAGTACAGCATCGGC |
| Cas9n-BE4max-R | Site-directed mutagenesis for removal of rAPOBEC1 from BE4max (reverse) | TCTTGTCGACTTTCCGCTTCTTCTTTGG |
| HAMMER-F | PCR HAMMER reporter linker region (forward) | CACTGCATACGACGATTCTGTG |
| HAMMER-R | PCR HAMMER reporter linker region (reverse); Reverse transcription of HAMMER mRNA | ATGTTCATCGAGTCCGACCC |
| HAMMER-seq | Sanger sequencing of HAMMER PCR products | TATTGTCGAGGGAGCTAAGA |

**Table S3. Viral ribonucleotide reductases.**

| Virus | Accession | Host Species | New World or Old World | Virus Family |
|---|---|---|---|---|
| HSV1 | YP_009137114 | Human | Old World | Alphaherpesvirinae |
| HCMV | YP_081503.1 | Human | Old World | Betaherpesvirinae |
| EBV | YP_001129452.1 | Human | Old World | Gammaherpesvirinae - lymphocryptovirus |
| KSHV | QLI54727.1 | Human | Old World | Gammaherpesvirinae - rhadinovirus |
| CalHV3 | NP_733909.1 | Marmoset | New World | Gammaherpesvirinae - lymphocryptovirus |
| SgHV1 | UNP64460.1 | Marmoset | New World | Gammaherpesvirinae - rhadinovirus |
| AtHV3 | NP_048032.1 | Atelinae | New World | Gammaherpesvirinae - rhadinovirus |
| SaHV2 | NP_040263.1 | Squirrel Monkey | New World | Gammaherpesvirinae - rhadinovirus |
| McHV13 | YP_010801332.1 | Macaque | Old World | Gammaherpesvirinae - lymphocryptovirus |
| McHV10 | YP_010084648.1 | Macaque | Old World | Gammaherpesvirinae - lymphocryptovirus |
| McHV4 | YP_067953.1 | Macaque | Old World | Gammaherpesvirinae - lymphocryptovirus |
| McHv5 | NP_570809.1 | Macaque | Old World | Gammaherpesvirinae - rhadinovirus |
| McHV8 | YP_010084428.1 | Macaque | Old World | Gammaherpesvirinae - rhadinovirus |

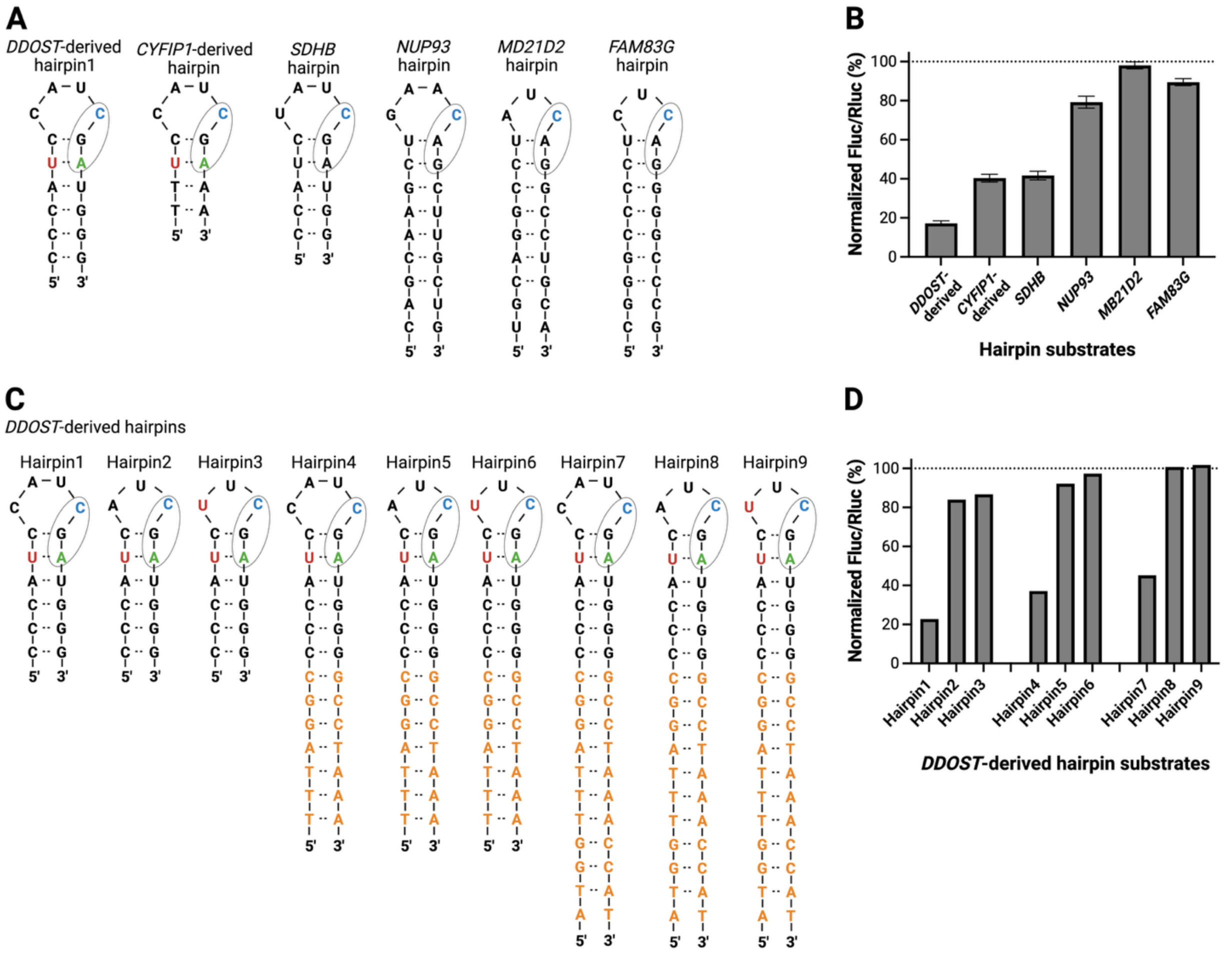


**Fig. S1. Additional hairpin substrates tested using HAMMER.**

**A**, Schematics of HAMMER reporter hairpin substrates derived from endogenous A3A-edited transcripts and engineered with variable loop residues and stem lengths. Target cytosines are highlighted in blue; bases modified from original sequences are colored in red (U) and green (A); stem bases extending from original sequences are shaded in orange.

**B**, Normalized firefly-to-renilla luminescence ratios for the indicated HAMMER reporters with different hairpin substrates co-expressed with human A3A in 293T cells [mean +/- SD of 2 biological reactions normalized to A3A-E72A control (dotted line at 100%)].

**C**, Schematics of DDOST1 hairpin1 and derivative hairpin substrates tested in the system described here. Target cytosines are highlighted in blue; bases modified from original sequences are colored in red (U) and green (A); stem bases extending from original sequences are shaded in orange.

**D**, Normalized firefly-to-renilla luminescence ratios for the indicated reporters with different DDOST-derived hairpin substrates co-expressed with human A3A in 293T cells [n=1 experiment normalized to A3A-E72A control (dotted line at 100%)].

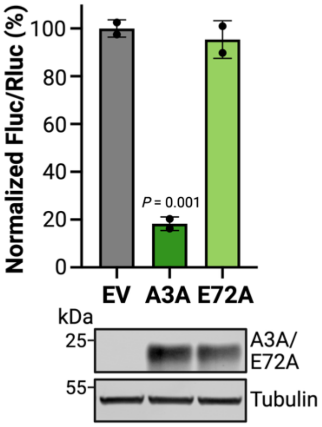


**Fig. S2. HAMMER reporter functionality in HeLa.**

Normalized firefly-to-renilla luminescence ratios of HeLa cells co-transfected with Hairpin1 reporter and either EV, A3A, or the catalytic mutant A3A-E72A (mean ± SD of 2 biological replicates). Immunoblots below confirm expression of A3A and A3A-E72A, with tubulin as a loading control.

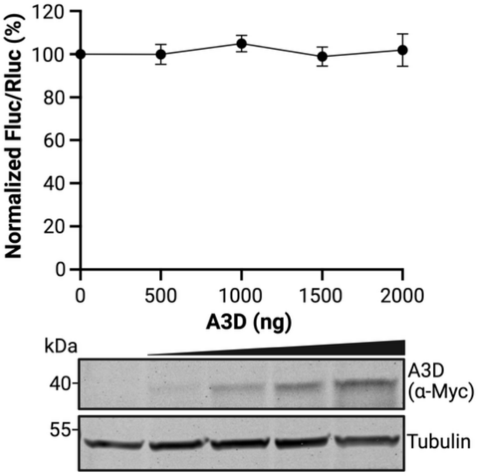


**Fig. S3. HAMMER is not a substrate for human A3D in 293T cells.**

Normalized firefly-to-renilla luminescence ratios of 293T cells co-transfected with Hairpin1 reporter and increasing amounts of human A3D (mean ± SD of 2 biological replicates). Immunoblots below confirm expression of A3D (anti-Myc) with tubulin as a loading control.

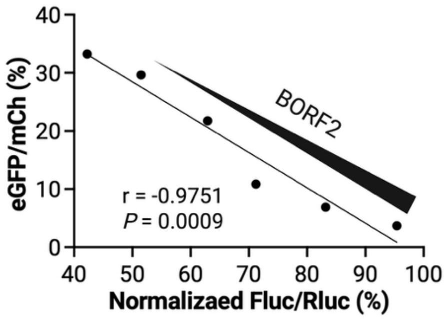


**Fig. S4. Relationship between DNA and RNA editing reporter activities**.

Correlation between HAMMER luminescence readout (x-axis) and AMBER fluorescence readout (y-axis) across increasing BORF2 expression levels in 293T cells co-expressing a fixed amount of A3A (200 ng) with each reporter. BORF2 was titrated using a two-fold dilution series starting from 400 ng plasmid. Pearson correlation coefficient (r) and corresponding *P*-value are indicated.
